# Cryo-electron tomography reveals the nanoscale architecture of the actin cortex in cellular blebs

**DOI:** 10.1101/2024.03.30.587413

**Authors:** Davide A.D. Cassani, Bruno Martins, Matthew B. Smith, Ohad Medalia, Ewa K. Paluch

**Author notes:** University Medical Center, Utrecht, the Netherlands (Matthew B. Smith). Corresponding authors: O.M. and E.K.P.

## Abstract

In animal cells, cellular deformations driving cytokinesis, migration, and epithelial constriction are driven by contractile tension in the actomyosin cortex, a thin network of actin and myosin underlying the plasma membrane. Cortical tension results from myosin-generated forces and as such, cortical myosin organization and dynamics have received significant attention. However, recent studies highlight that alongside myosin motor activity, the organization of the cortical actin network is a key regulator of tension. Yet, very little is known about the structural arrangement of cortical actin filaments. This is mostly due to the small thickness and high density of the cortex, which makes the visualization of cortical actin filaments challenging. Here, we use cryo-electron tomography (cryo-ET) to unveil the structural organization of cortical actin. As a model, we use isolated cellular blebs, which assemble an actin cortex comparable to the cortex of entire cells, but are small enough to be amenable to cryo-ET. We find that the bleb actin cortex is mostly composed of short and straight actin filaments. We then characterize cortex structural parameters, including the density of potential cross-linking and membrane attachment points. Our study unveils the nanoscale three-dimensional organization of the cortical actin network in cellular blebs. As such, it provides a quantitative framework for models of cortical tension generation, and will help understanding the nanoscale basis of cell surface contractions.

## Introduction

Key biological processes, including cell migration, division and tissue morphogenesis, are mediated by changes in cell shape. In animal cells, shape changes are primarily controlled by the cellular cortex, a thin actomyosin network supporting the plasma membrane (Salbreux et al., 2012). The cortex is under myosin-generated contractile tension, and tension gradients drive cellular deformations (Chugh and Paluch, 2018; Heisenberg and Bellaiche, 2013; Levayer and Lecuit, 2011). Understanding cortex tension regulation is thus essential to understanding cellular morphogenesis.

Cortical tension is often equated to the level of active myosin II at the cortex (Bergert et al., 2015; Fierling et al., 2022; Mayer et al., 2010). However, recent studies have highlighted that the spatial organization of cortical actin filaments is equally important (reviewed in (Koenderink and Paluch, 2018; Murrell et al., 2015)). For instance, depletion of proteins controlling actin filament length or cross-linking has been shown to affect cortical tension (Chugh et al., 2017; Ding et al., 2017; Toyoda et al., 2017). This is consistent with *in vitro* studies and modeling, which have shown that actin organization, and in particular the level of actin network connectivity, are key contractile tension regulators (Chugh et al., 2017; Ennomani et al., 2016).

Despite its importance, the organization of actin filaments at the cortex remains poorly understood due to the small thickness and high density of the cortical network (Chugh and Paluch, 2018). Scanning electron microscopy (SEM) and atomic force microscopy (AFM) have been used to map the surface of the cortex, unveiling a dense filament network with a meshsize typically smaller than 50-100 nm in tissue culture cells (Bovellan et al., 2014; Chugh et al., 2017; Eghiaian et al., 2015; Kronlage et al., 2015). Electron tomography of the cytoplasmic side of the cortical network remaining at the plasma membrane after the rest of the cell is torn off, revealed meshsizes between 50 and 200 nm, depending on the cell type (Morone et al., 2006); however, it is unclear to what extent the cortex is preserved during the membrane rip off process. Altogether, these techniques only probe one surface of the cortex and much less is known about the three-dimensional (3D) architecture of the network. Sub-resolution image analysis and super-resolution microscopy techniques indicate that the thickness of the cortical actin layer is 100-400 nm (Chugh et al., 2017; Clark et al., 2013; Clausen et al., 2017), in line with physical measurements where the cortex is “pinched” between two magnetic beads (Laplaud et al., 2021). Due to this very low thickness, coupled to the high density of the cortical network, resolving cortical actin organization in the direction transversal to the plasma membrane has so far not been possible using optical microscopy, including super-resolution techniques. As a result, the 3D organization of actin filaments at the cortex remains essentially a black box. This has seriously limited the scope and realism of theoretical models of cortical tension generation, which have to rely on assumptions on cortical actin architecture that remain unverified (reviewed in (Koenderink and Paluch, 2018)).

Cryo-electron tomography (cryo-ET) is increasingly the method of choice to explore intracellular 3D architecture (Chakraborty et al., 2020; Chung et al., 2022; Schuller et al., 2021; Zhang et al., 2023). However, to generate good quality tomograms, Cryo-ET requires samples thinner than ∼ 1 μm (Lucic et al., 2005; Weber et al., 2019), while the cortex is usually most prominent in thick cellular regions, far from the substrate and from adhesive regions. A recent study has used focused-ion-beam (FIB) milling to generate thin equatorial lamellae across pairs of neighboring cells, revealing cortical actin filaments arranged mostly parallel to the plasma membrane, with prominent actin bundles (Lembo et al., 2023). This arrangement contrasts with SEM images of the actin cortex at the surface of rounded cells, which suggest a more isotropic network (Bovellan et al., 2014; Chugh et al., 2017). Overall, the 3D architecture of the actin cortex at free membranes in rounded parts of the cell, where it is most prominent, remains poorly understood.

Here, we used small isolated cellular blebs, which re-assemble a functional cortex similar to the cortex of intact cells (Biro et al., 2013; Bovellan et al., 2014), as a simplified model of the cortex for cryo-ET investigation. We optimized the bleb isolation procedure to obtain large numbers of blebs < 1 μm in diameter, amenable to cryo-ET. We characterized the actin cortex in these small blebs using super-resolution microscopy. We then analyzed small isolated blebs using cryo-ET. We could show that the blebs contain a dense network of actin filaments, and characterized filament properties and density, as well as the density of potential sites of cross-linking, branching and attachment to the plasma membrane. Together, our analysis unveils that the actin cortex in isolated blebs comprises mostly short, straight filaments, and provides a quantitative description of the structural organization of the cortical network.

## Results

### Preparation of cortex-enriched blebs for cryo-electron tomography

We first established a protocol for the isolation of cortex-rich cellular blebs for cryo-ET (Fig. 1A,B). We used HeLa cells, where cortex composition and mechanics have been extensively characterized (Chugh et al., 2017; Ramanathan et al., 2015; Serres et al., 2020; Toyoda et al., 2017; Truong Quang et al., 2021), and adapted a bleb isolation method we previously developed (Biro et al., 2013; Vadnjal et al., 2022). Strong blebbing was induced by short treatment with the actin depolymerizing drug latrunculin B (Fig. 1A *(i,ii)*) and blebs were detached from the cell body by mechanical agitation (Fig. 1A *(iii, iv)*). Latrunculin was then washed out and an ATP regeneration system was added to the blebs (see Methods) to facilitate cortex reassembly (Biro et al., 2013). Finally, in order to enrich for blebs small enough to be amenable to cryoET, the supernatant was filtered through a 5 µm-pore-size filter and blebs were collected by centrifugation (Fig. 1A *(v)*, B). The filtering allowed to separate small blebs from larger blebs, debris and detached cells (Fig. 1B). This protocol yielded large numbers of blebs smaller than 1 µm in diameter (Fig. 1B, arrows in right image), a size amenable to cryo-ET imaging.

**Figure 1:**
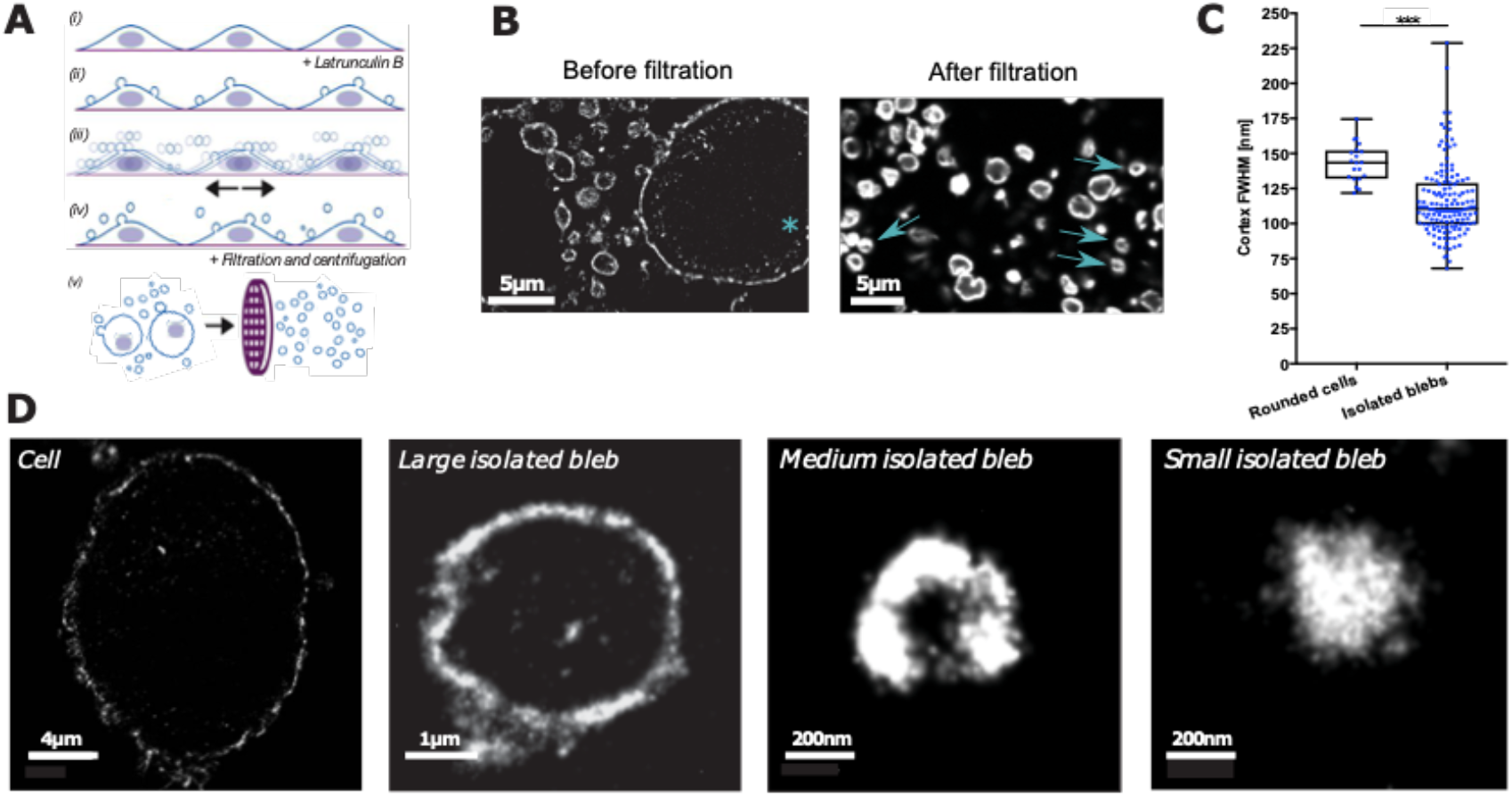
Isolation and characterization of cortex-enriched blebs. (A) Schematic of the bleb isolation protocol: addition of Latrunculin B (*i*) leads to intensive blebbing (*ii*), blebs are detached from cells by mechanical agitation (*iii*, *iv*), and small blebs are enriched by filtration of the supernatant (*v*). (B) Representative confocal images of the supernatant obtained as described in (A) before (left) and after (right) filtration. Actin was labeled with phalloidin. Before filtration the supernatant comprises blebs of a variety of sizes, as well as detached cells (marked with *). After filtration only small blebs are present; arrows point to blebs of sizes amenable to cryo-ET. (C) Actin cortex FWHM, as an estimate of cortex thickness, measured in dSTORM images of rounded-up cells and in isolated blebs. Mean values (± standard deviation): 116 ± 26 nm for n=144 blebs vs. 143 ± 14 for n = 20 cells, 3 independent experiments. Boxplots indicate median, 25^th^ and 75^th^ percentile. Statistics: given the difference in sample sizes, hypothesis testing was performed using bootstrapping, p-value = 0.0002. (D) Representative dSTORM images of actin in a rounded cell (left) and in isolated blebs of different sizes.

We then verified that the cortex reassembled in isolated blebs was comparable to that of intact cells, as suggested by our previous analysis (Biro et al., 2013). Confocal imaging of actin showed the presence of a cortical actin layer in isolated blebs (Fig. 1B). However, the cortical layer is under the resolution of classical light microscopy (Clark et al., 2013) and thus confocal microscopy-based characterization is limited. Therefore, we turned to super-resolution microscopy and compared actin cortices in isolated blebs and rounded HeLa cells, where the cortex is prominent (Chugh et al., 2017), using direct Stochastic Optical Reconstruction Microscopy (Truong Quang et al., 2021) (dSTORM; Fig. 1C,D). We observed a clear cortical layer, comparable to the cortex of intact cells, in most isolated blebs larger than ∼ 500 nm (Fig. 1D). Most blebs smaller than ∼ 500 nm did not display a well-defined cortex but rather appeared filled with actin (Fig. 1D, right); this is likely because the cortex fills the entire volume of such small blebs and thus cortex-free cytoplasm cannot be visualized. We then measured the full width at half maximum (FWHM) of the cortical actin layer, as a readout of cortex thickness (Serres et al., 2020; Truong Quang et al., 2021), in isolated blebs with a clear cortex and in rounded cells. We found that cortical thickness was of the same order of magnitude, though slightly smaller, in isolated blebs compared to entire cells (Figs. 1C and S1). Taken together, these data indicate that our protocol allows for the isolation of blebs small enough for cryo-ET, containing an actin cortex of thickness comparable to the cortex of entire cells.

### Visualization of actin filaments in isolated blebs by cryo-ET

We then applied cryo-ET to isolated blebs. Isolated blebs were blotted onto carbon-coated grids and vitrified, and tilt series of electron microscopy images of small blebs were acquired and used for three-dimensional reconstruction (Fig. 2A,B, Supplementary Movie 1). Back-projected tomograms revealed clear filamentous structures (Fig. 2B) with diameters consistent with the diameter of an actin filament (∼7.5 nm, Fig. 2C). Filaments manually distinguishable by visual inspection are mostly those oriented around the x-y plane rather than parallel to the electron beam. To detect and analyze filament organization in 3D, we used an algorithm developed for actin segmentation in tomograms (Rigort et al., 2012) (Fig. 2D, Supplementary Movies 2 and 3). Due to the missing wedge effect, the top and bottom of a vesicle-like structure can hardly be detected in cryo-tomograms (Grimm et al., 1997). Therefore, we segmented and analyzed only 100-200 nm tomogram sections around the mid-planes of blebs, in which both actin and the membrane are well detectable. Within these tomographic sections, the algorithm successfully detected actin filaments with various orientations (Fig. 2D,E). This approach allowed us to segment dense filamentous networks in 10 purified blebs (Fig. S2).

**Figure 2:**
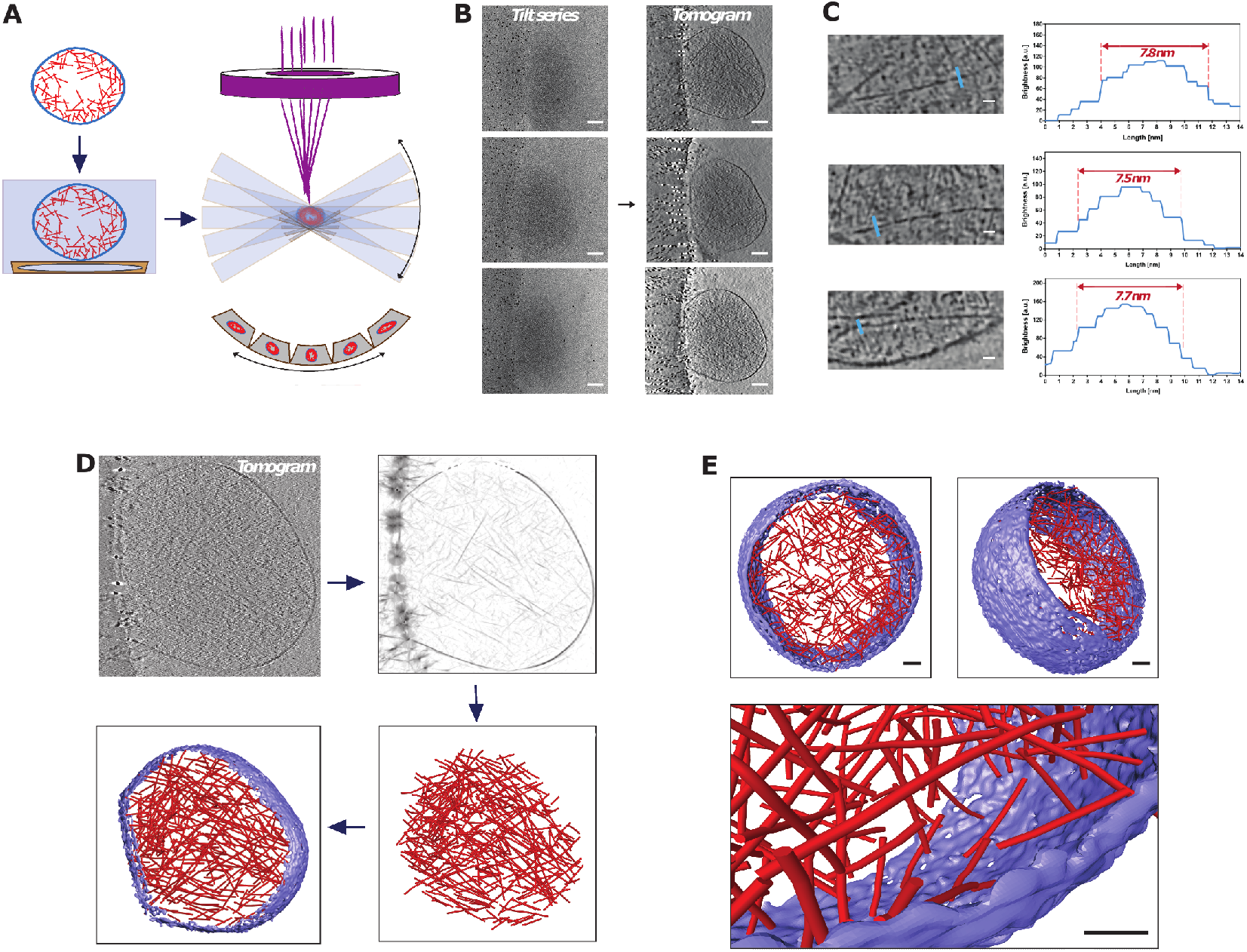
Cryo-ET imaging of isolated blebs. (A) Schematic illustration of the cryo-electron tomogram acquisition procedure for isolated blebs. (B) Examples of tilted projection images and respective reconstructed tomograms. Scale bars, 100 nm. (C) Left: filamentous structures from tomograms of isolated blebs. Right: intensity profiles along blue lines highlighted in images on the left. The width of the intensity profiles, 7.5-8 nm, is consistent with the diameter of an actin filament. Scale bars, 20 nm. (D) Schematic of the automatic 3D segmentation of filaments (red) and plasma membrane (blue). (E) Example views of a segmented section of an isolated bleb. Scale bars, 50 nm. All the blebs segmented are displayed in Fig. S2. The bleb displayed in (E) is the same as in Fig. S2D.

To further test whether the filamentous structures identified were indeed actin filaments, we treated isolated blebs with the F-actin depolymerizing agents cytochalasin D, which caps filament barbed ends and prevents polymerization, and latrunculin A, which sequesters actin monomers (thereafter CDLA treatment). Overall, CDLA treatment led to significant reduction in actin amounts at the cortex of isolated blebs, with most of the F-actin collapsed into aggregates (Fig. 3A), as previously reported upon treatment with actin depolymerizing drugs (Wakatsuki et al., 2001). Cryo-ET imaging showed a reduction in filamentous networks in most CDLA-treated blebs (Figs. 3B,C and S3). A few of the CDLA-treated blebs appeared to preserve a dense filamentous network (Fig. S3A blebs b,e,f,h), and reduction in filament density was generally weaker in smaller blebs (Fig. S3B). This suggests that actin filaments may be stabilized against CDLA action in blebs below a certain size. Nonetheless, in the majority of the 16 blebs imaged, CDLA treatment led to a strong reduction of the filamentous network (Figs. 3 and S3). Together with the diameter of the observed filaments (Fig. 2C), these findings support the conclusion that cryo-ET imaging reveals actin filament organization in isolated blebs.

**Figure 3:**
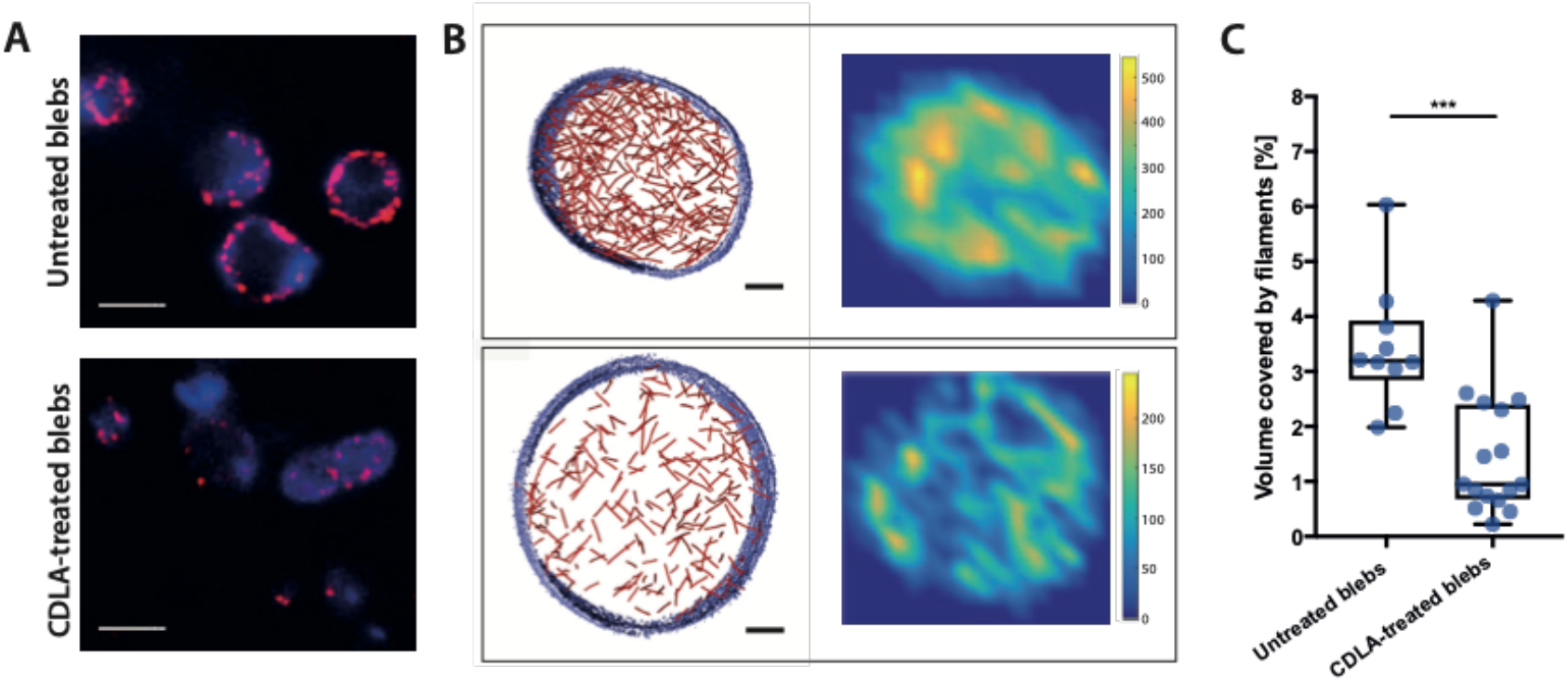
Actin depolymerizing agents disrupt filament networks in isolated blebs. (A) Correlative dSTORM and confocal images of untreated (top) or CDLA-treated (bottom) isolated blebs. Membrane is labeled in blue (CellMask Green, confocal) and F-actin in red (phalloidin Alexa Fluor 647 nm, dSTORM). Scale bars, 1 μm. (B) Segmented tomograms (left) and filament density maps (right, scales indicate intensity in a.u.) in an untreated bleb (top, same bleb as in Fig. S2E) and a CDLA-treated treated bleb (bottom; same bleb as bleb (m) in Fig. S3A). All the CDLA-treated blebs segmented are displayed in Fig. S3A. Scale bars, 100 nm. (C) Quantification of volume occupied by filaments in the tomograms of untreated (n=10) and CDLA-treated (n=16) blebs; 2 independent experiments. Boxplots: median, 25^th^ and 75^th^ percentile. Statistics: Student’s t-test, p<0.001.

Notably, the actin networks observed did not always form a clear cortical layer (Fig. S2). Our dSTORM measurements show that the half-width of the bleb actin cortex is ∼120 nm (Figs. 1C and S1B,C), suggesting actin filaments extend over 200 nm from the plasma membrane. As the radii of the blebs we could analyze by cryo-ET (Fig. S2) were between ∼170 and ∼530 nm, cortical actin would be expected to almost entirely fill these small blebs, accounting for the lack of a clearly-defined actin-free central region. To further assess this, we plotted filament density maps in the segmented blebs (Figs. 3B and S2). In several blebs, density maps suggest reduced filament density towards the bleb center (Fig. S2C,D,E,I,J). Taken together, our observations indicate that the filamentous network observed by cryo-ET in isolated blebs is the F-actin cortex.

### Quantitative analysis of actin cortex architecture in isolated blebs

We then analyzed the spatial architecture of the bleb F-actin cortex. The density of actin filaments in isolated blebs was 8,107 (6,608 – 10,562, median (25° - 75° percentile)) filaments / μm^3^ (Fig. 4A), corresponding to 3.19 % (3.07 – 3.71, median (25° - 75° percentile)) of the bleb volume being occupied by actin filaments (see Methods for details on calculations). Actin filament density appeared independent of bleb size (Fig. S3B). Using the total segmented filament length, we estimated that the concentration of F-actin in isolated blebs was ∼380 µM. Filament length is challenging to determine precisely in cryo-electron tomograms because local loss of contrast may be interpreted by the software as a filament edge, leading to mis-identification of two portions of an individual filament as two filaments. Furthermore, many filaments extend beyond the section that was imaged and their lengths can therefore not be measured. Nonetheless, we analyzed the lengths of the segmented filaments that were entirely contained within our tomogram slices (Fig. 4B) and found that most were very short with 84 % shorter than 100 nm. Notably, given that our analysis software only detects filaments longer than 54 nm (see Methods), the mean filament length we report is likely an over-estimate. The median diameter of the blebs investigated was 627 nm (Fig. S2), suggesting that filament length is not simply limited by bleb size. Finally, we quantified filament straightness (Swulius et al., 2018) (Figs. 4C and S4A). As also apparent from visual inspection of the segmented F-actin networks (Fig. S2), the actin filaments in isolated blebs displayed almost no curvature, with 71% of the filaments displaying a straightness higher than 0.99 (Fig. 4C). This is consistent with the fact that the length of the actin filaments in isolated blebs is very small compared to the persistence length of F-actin (∼10 µm, (Isambert et al., 1995)). Taken together, our analysis indicates that the bleb actin cortex comprises a dense network of relatively short, straight filaments.

**Figure 4:**
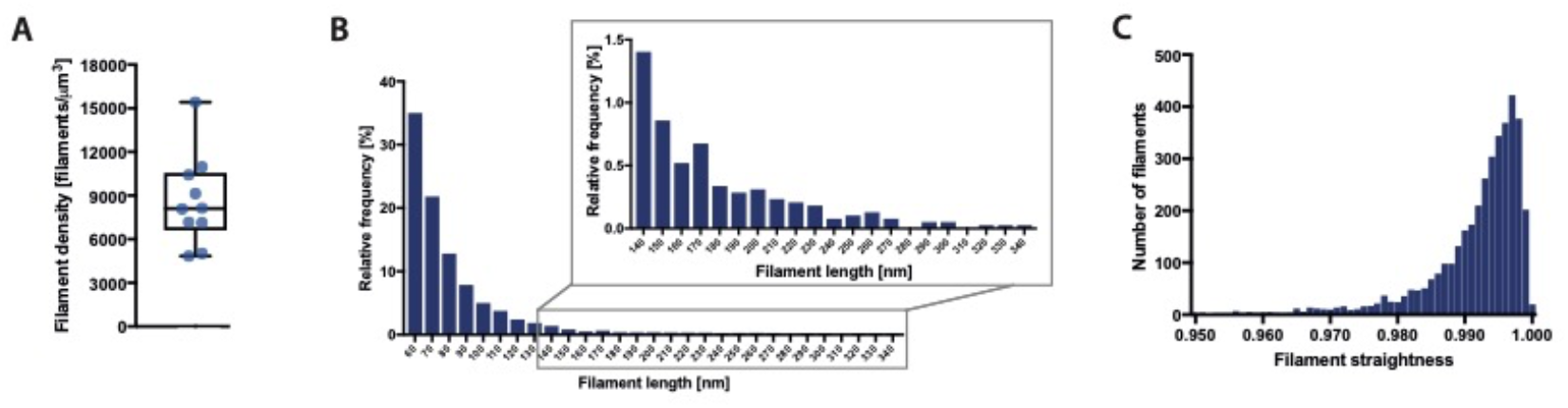
Analysis of actin filaments in the bleb cortex. (A) Filament density in isolated blebs; n = 10 blebs in 2 independent experiments. Boxplot: median, 25^th^ and 75^th^ percentile. (B) Filament length distribution in isolated blebs. Note that the software only identifies filaments longer than 54 nm (see Methods). (C) Distribution of filament straightness, defined as the Euclidean distance between the filament ends divided by the contour length of the filament. For B and C: n= 3829 filaments in a total of 10 blebs from 2 independent experiments.

We then investigated potential filament-filament interactions in the segmented F-actin networks. For this, we analyzed network organization using custom-written software. Segmented filaments were partitioned into filament heads (2 nm caps at the filament ends) and filament body (remaining section between the two heads), and we searched for potential crossing and branching points where filaments came close enough to each other so that cross-linkers or branching proteins could connect two filaments (Fig. 5A).

**Figure 5:**
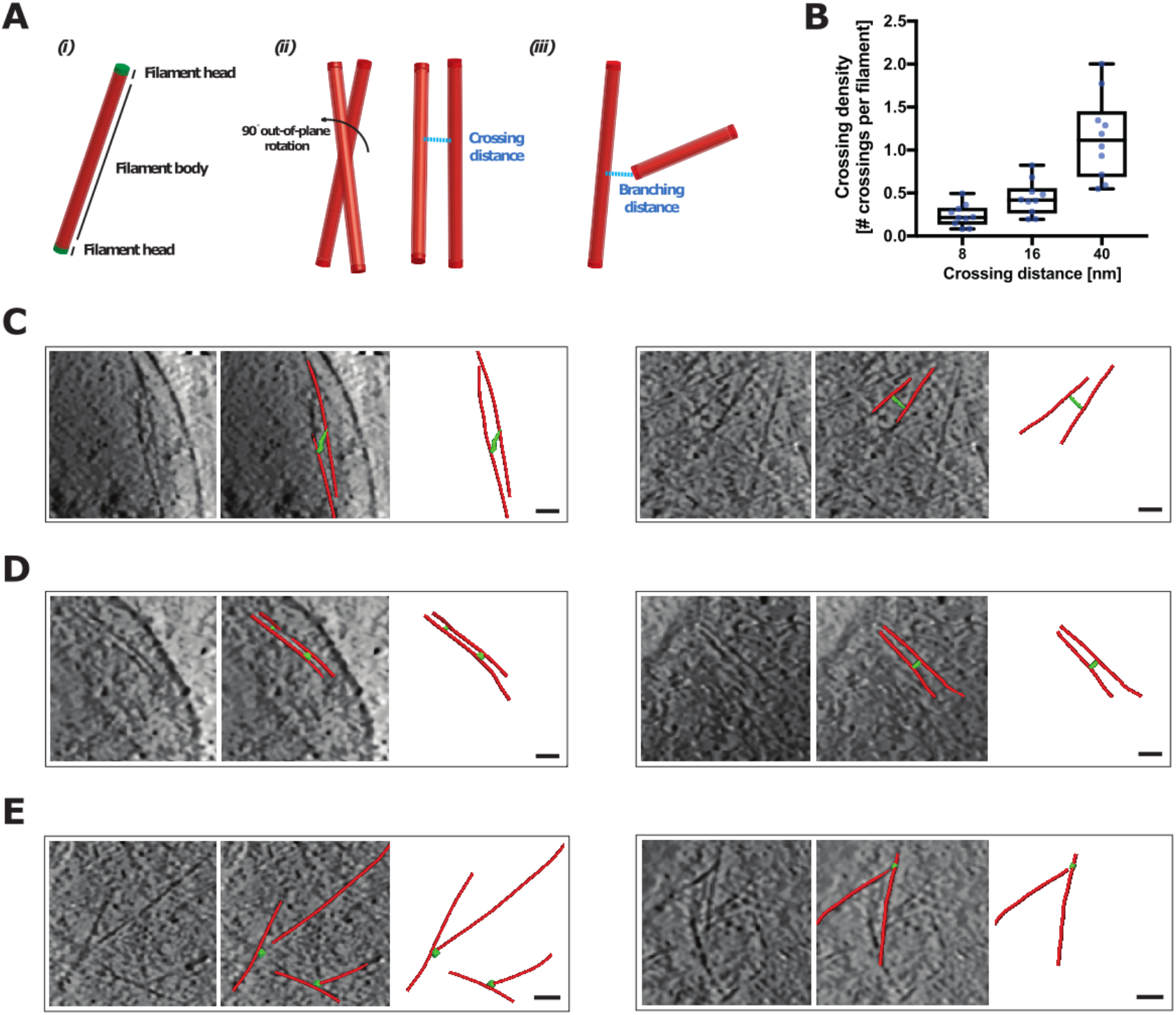
Filament crossing and branching. (A) Graphical representation of an actin filament and of possible filaments configurations: *(i)* single filament with heads highlighted in green and filament body in red, *(ii)* filament crossing with crossing distance (blue dashed line), *(iii)* filament branching with branching distance (blue dashed line). (B) Crossing density for different crossing distances: contact crossings at 8 nm, short- and medium-distance crossings at 16 (± 3) nm and 40 (± 3) nm, respectively, in n=10 isolated blebs. Boxplots: median, 25^th^ and 75^th^ percentile. (C) Examples of long electron-dense structures (green) appearing to connect two actin filaments (red). (D) Examples of short electron-dense structures (green) appearing to connect two actin filaments (red). (E) Examples of filament branching points (highlighted in green). (D-E) Scale bars, 50 nm.

A filament crossing point was recorded when the bodies of two filaments crossed each other at a defined distance (“crossing distance”) (Fig. 5A-D). Notably, actin crosslinkers greatly vary in size. A variety of different cross-linkers have been identified at the cortex (Biro et al., 2013), including fascin, which cross-links actin filaments 8 nm apart, and alpha-actinin, which is ∼ 35 nm in size (Winkelman et al., 2016). Taking into account that the crossing distance is measured from the centers of the actin filaments, which have a radius of ∼ 4 nm, we thus quantified crossing density per filament at 16 nm (corresponding to a distance between the surfaces of the two filaments of ∼ 8 nm) and at 40 nm (corresponding to a distance between the surfaces of the two filaments of ∼ 32 nm, Fig. 5B). We found that, crossing points at 16 nm (± 3 nm), potential cross-linking sites for short cross-linkers, were relatively rare, with on average 1 crossing every 2 filaments (Fig. 5B). We observed ∼ 1 crossing per filament at 40 ± 3 nm (Fig. 5B). In some cases, we could observe electron-dense structures connecting neighboring filaments that may be cross-linkers (Fig. 5 C, D). We also found that about 1 in 4 filaments crossed another filament at an 8 nm distance (Fig. 5B), corresponding to actin filaments practically touching each other. Uncertainty in the segmentation of filament extremities prevented a precise quantification of branching points, however some clear branches could be observed in the network (Fig. 5E).

Finally, we analyzed possible points of contact between actin and the plasma membrane (Fig. 6). We quantified the density of filament-membrane contact points, defined as locations where a filament head was in direct contact with the plasma membrane (Fig. 6A,C). We found ∼120 contact points per µm^2^ of plasma membrane (Fig. 6B). This number did not appear to depend on the size of the bleb (Fig. S4B). Furthermore, we occasionally observed electron-dense structures connecting the body of the filaments to the plasma membrane (Fig. 6D). These could be members of the ezrin-radixin-moesin family or myosin 1 motors, which are known to connect actin to the plasma membrane (Chugh and Paluch, 2018). Taken together, our analysis provides a quantitative description of the nanoscale architecture of the actin cortex in isolated blebs (Table 1).

**Figure 6:**
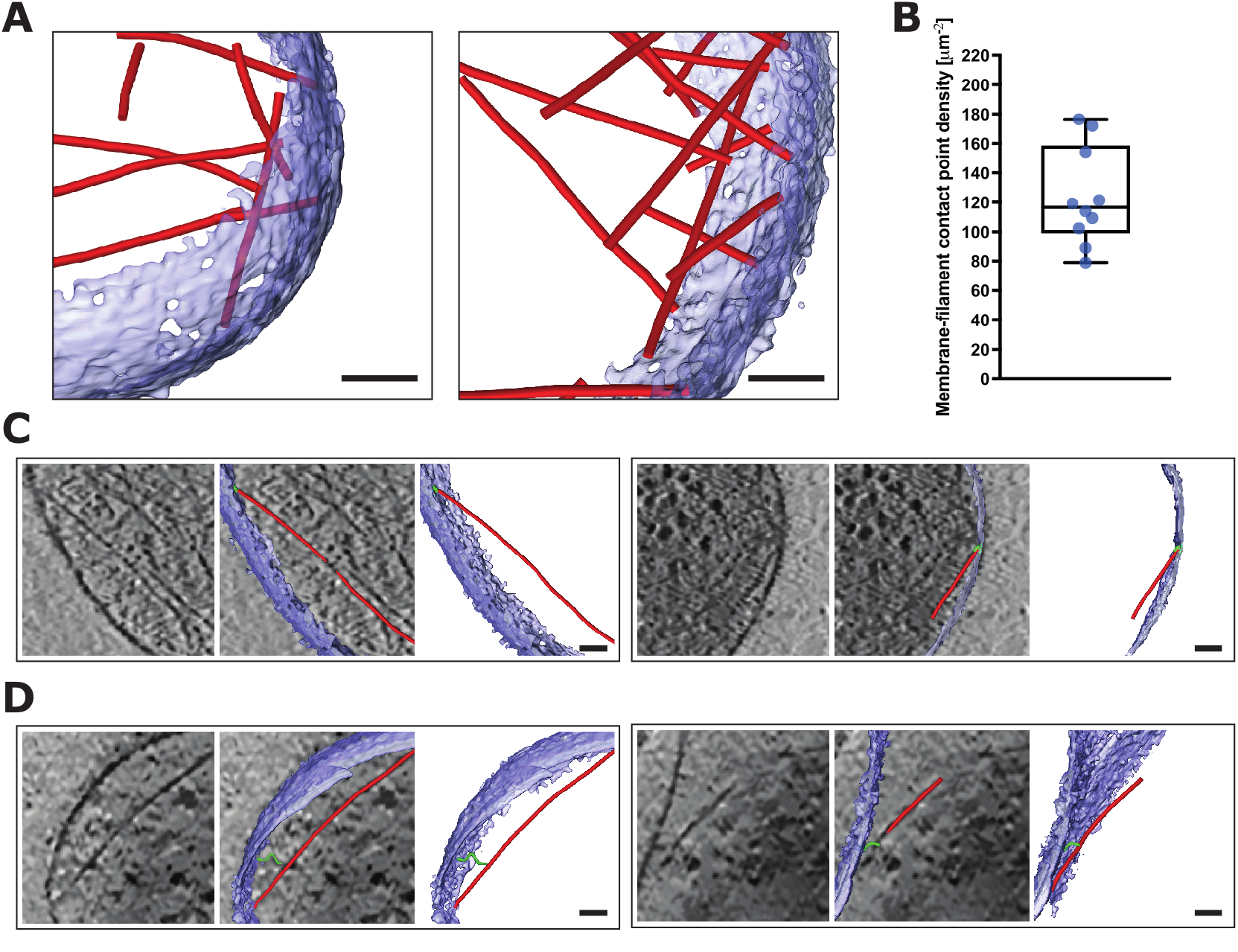
Connections between filaments and the plasma membrane. (A) Examples of actin filaments (red) in contact with the plasma membrane (blue) in isolated blebs. (B) Density of filament-membrane contact points in n=10 isolated blebs. Boxplot: median, 25^th^ and 75^th^ percentile. (C) Example tomograms where a filament end appears in direct contact with the plasma membrane. The filament end in contact with membrane is marked in green. Quantified in B. (D) Example tomograms where an electron-dense structure (green) appears to connect a filaments body (red) to the plasma membrane (blue). Scale bars, 50 nm.

**Table 1:** Parameters describing the nanoscale architecture of the actin cortex in isolated blebs. Values are medians (25^th^ – 75^th^ percentile)

| Parameter | Definition | Unit | Value |
| --- | --- | --- | --- |
| Filament number | Number of filaments | # | 397 (215-498) |
| Filament density | Number of filaments per unit volume | #/ $\mu\text{m}^3$ | 8107 (6608 - 10562) |
| F-actin molarity | Molarity of polymerized actin (estimated from total filament length) | $\mu\text{M}$ | 380 (339 - 468) |
| Filament length | Estimated mean filament length | nm | 76.2 (61.8 - 86.1) |
| Straightness | Displacement length along actin filament divided by contour length | - | 0.993 (0.989 - 0.996) |
| Density of contact crossings (8 nm crossing distance) | Density of crossing locations where two filaments touch each other | #/ 100 nm | 0.28 (0.15 - 0.41) |
| Density of short-distance crossings (16 nm crossing distance) | Density of crossing locations with 8 nm between the outer surface of two filaments | #/ 100 nm | 0.54 (0.29 - 0.69) |
| Density of medium-distance crossings (40 nm crossing distance) | Density of crossing locations with 32 nm between the outer surface of two filaments | #/ 100 nm | 1.35 (0.78 - 1.82) |
| Density of membrane-cortex contact points | Density of filaments in direct contact with the bleb membrane | #/ $\mu\text{m}^2$ | 116 (99 - 158) |

## Discussion

In this study, we isolated cellular blebs to investigate the structural organization of the sub-membranous actin cortex using cryo-ET. A similar approach has been used for the study of the adhesion machinery in platelets, where thin platelet-derived microparticles were used to obtain samples amenable to cryo-ET (Tamir et al., 2016). Previous scanning electron microscopy observations indicate that the bleb cortex is similar to the cortex of entire cells, and can display contractile behavior upon addition of ATP (Biro et al., 2013). Our dSTORM characterization further shows that the thickness of the bleb cortex is on the same order as cortex thickness in intact cells (Figs. 1C and S1). Taken together, these observations strongly suggest that isolated blebs are a good model system to investigate cortical actin organization by cryo-ET.

Our cryo-ET analysis reveals a dense network of short straight actin filaments in isolated blebs and yields important parameters describing the network architecture. We found that actin filaments occupied around ∼3.2 % of the volume of isolated blebs, corresponding to an F-actin concentration of around 380 µM. This is comparable to the F-actin concentration in lamellipodia estimated by cryo-ET analysis to be ∼500 µM (Koestler et al., 2009). Interestingly, our analysis revealed that actin filaments in the cortex of isolated blebs are mostly straight and shorter than 100 nm (Fig. 4). This is shorter than the filaments observed in tomograms of stress fibers or membrane protrusions (Rigort et al., 2012) (average lengths of 320 nm and 168 nm, respectively), or at the core of podosomes (average length 111 nm (Jasnin et al., 2022)), and than the actin filaments imaged by FIB milling/cryo-ET at the membrane of adjacent rounded fibroblasts (Lembo et al., 2023). It is possible that the small size and high membrane curvature of the blebs we could analyze using cryo-ET limits actin filament length. It is also possible that cortex organization is context-dependent. Further studies, using FIB milling to image the cortex in a variety of contexts will be required to uncover how properties like local membrane curvature or outside constraints affect cortex organization.

Even though our cryo-ET images revealed some electron-dense structures that could be actin cross-linkers (Fig. 5 C-E), the resolution of the tomograms was not sufficient to systematically identify cross-linkers. Instead, we characterized locations where two actin filaments crossed each other at a distance compatible with the binding of cross-linkers of different sizes. We found 1 crossing every 2 actin filaments compatible with short cross-linkers, such as fascin (6 nm long) or fimbrin (10 nm long) (Winkelman et al., 2016), and 1 crossing per filament at a distance that could accommodate medium-sized crosslinkers like alpha-actinin (35 nm long, (Winkelman et al., 2016)). Of note, any short-distance crossing could be compatible with the binding of a longer cross-linker, provided flexibility in attachment angle between the cross-linker and actin. We also found that 1 filament in 4 was in direct contact with another filament, which could accommodate any size of cross-linker. Taken together, our data suggest that the bleb cortical network could accommodate 1 to 2 cross-links of the size of alpha-actinin or smaller, per actin filament. We also observed clear branches in the network, but uncertainties on filament extremities precluded quantification of branching density. Finally, we identified about 120 µm^-2^ contact points between actin filaments and the plasma membrane, corresponding to about 100 nm distance between actin-membrane contacts. This number is an under-estimate, as it does not take into account locations where an actin filament is attached to the membrane by a membrane-actin linker, without direct contact between the filament and the membrane.

Our quantitative characterization of the architecture of the actin cortex will be of direct relevance for theoretical models of cortex contractility. A variety of computational models investigating how network properties affect contractile tension generation have been developed (reviewed in (Koenderink and Paluch, 2018; Murrell et al., 2015)). They mostly rely on parameters derived from in vitro studies, or explore parameter spaces based on assumptions of what the cortical actin network might look like. Interestingly, several recent models have highlighted that connectivity, or the number of cross-links per filament, is a key regulator of contractile tension, with optimal tension corresponding to an intermediate level of network connectivity (Belmonte et al., 2017; Chugh et al., 2017; Ennomani et al., 2016). A computational model suggests that optimal connectivity could be achieved at ∼4 cross-links per actin filament (Ennomani et al., 2016). Our study identified 1 to 2 crossing points at short to medium distances per actin filament, suggesting that longer cross-linkers, such as filamin (>100 nm between actin binding sites (Nakamura et al., 2011)), might be required for optimal contractility. However, the exact value of optimal connectivity is expected to depend on other parameters, such as motor density, filament straightness, and possibly the degree of branching (Belmonte et al., 2017; Ennomani et al., 2016). Future studies with enhanced resolution will be required to precisely quantify these parameters. It will also be interesting to develop further cryo-ET approaches to probe how cortical architecture changes during the generation of contractile tension gradients, bringing further insight into how nanoscale molecular interactions control cellular morphogenesis.

## Acknowledgements

We thank M. Serres and N. Vadnjal for technical support with the bleb isolation protocol, B. A. Truong Quang for help with the dSTORM experiments, and the LMCB electron microscopy facility, in particular J. Burden, for technical advice. We thank G. Charras for advice on the project. This work was funded through grants from the European Research Council (Consolidator grant 820188-NanoMechShape to E.K.P. and 810057 to O.M.), the Medical Research Council UK (MRC Programme Award MC_UU_12018/5 to E.K.P.), and the Human Frontier Science Program (Young Investigator Grant to E.K.P.) and the Schweizerischer Nationalfonds (310030_207453 to O.M.).

## Author contributions

D.A.D.C., O.M. and E.K.P. designed the research; D.A.D.C. carried out all the experiments and analyzed the data; B.M. helped with the cryo-EM experiments and tomogram analysis; M.B.S. developed software for actin network architecture analysis; D.A.D.C., O.M. and E.K.P. wrote the paper.

## Materials and methods

### Cell culture and cell synchronization

HeLa TDS cells were a gift from the MPI-CBG Technology Development Studio (TDS, Dresden, Germany). Cells were cultured in DMEM (Gibco by Life Technologies) with 1% penicillin/streptomycin, 1% glutamine, and 10% fetal bovine serum (Gibco by Life Technologies). Cell culture was carried out in polystyrene T75 culture flasks (Thermo Scientific). Cells were routinely tested for mycoplasma. To synchronize cells in interphase (G1/S-phase), cells were incubated overnight (∼16 h) with 2 mM thymidine (Sigma-Aldrich).

### Bleb isolation and CDLA treatment

Cells were grown to confluence in T75 tissue flasks (Thermo scientific) and synchronized in interphase. Prior to the experiment, the flask was gently manually shaken to detach debris and dead cells, and the medium was removed and replaced with 4 mL of fresh medium supplemented with 2.1 μM Latrunculin B (Sigma-Aldrich). Flasks were then taped to an Eppendorf Thermomixer Compact and shaken for 15 min at 750 rpm at room temperature. The supernatant containing isolated blebs and some detached cells was collected, resuspended in an extra 6 mL of medium, and spun down using an Eppendorf 5804R centrifuge for 5 minutes at 4400 rcf. The pellet was resuspended in 2 mL of low calcium intracellular buffer (IB, 10 mM NaCl, 280 mM K-Glutamate, 14 mM MgSO_4_, 13.3 mM CaCl_2_, 20.4 mM K-EGTA, 40 mM K-HEPES, pH 7.2). The 2 mL suspension was loaded into a 2 mL syringe and filtered through a Minisart syringe end filter (pore size 5.0 μm). To avoid filter clogging, bleb suspensions from different flasks were combined only after the filtration step. Three flasks were used for each experiment. After filtration, blebs were resuspended and incubated for 30 minutes in IB containing 2% of an exogenous ATP regeneration system (Energy mix: 1 mM UTP, 1 mM ATP, 6.1 μM creatine phosphokinase, 10 mM creatine phosphate in IB) and 50 μg/ml of *Staphylococcus aureus* α-toxin (Hemolysin, Sigma-Aldrich) to permeabilize the bleb membrane and allow for ATP and UTP to penetrate the blebs. All the reagents for IB buffer and α-toxin were purchased from Sigma-Aldrich, reagents for energy mix were purchased from Merck. For the CDLA treatment, isolated blebs were resuspended in 2.1 μM Latrunculin B, 0.5 μM Latrunculin A and 3.9 μM Cytochalasin D (all from Sigma Aldrich) for 15 minutes prior to fixation or transfer onto cryoET grids.

### Isolated blebs fixation for light and super resolution microscopy

Isolated blebs prepared as described above were spun onto glass coverslips. Fixation and membrane permeabilization were carried out with 4% PFA and 0.2% TritonX for 7 minutes, followed by 14 minutes of further fixation with 4% PFA alone. After 3 washes in IB, isolated bleb cortices were incubated with 0.5% phalloidin 647 nm (Life-Technologies) for 20 minutes. After 3 washes, coverslips were mounted onto glass slides and sealed with nail polish. For additional membrane labeling, after incubation with phalloidin 647 nm, coverslips were washed and incubated with 0.04% Cell Mask green (Life-Technologies) in IB for 10 minutes.

### dSTORM and confocal imaging

dSTORM imaging was performed on a Nikon N-STORM inverted optical microscope. Images were acquired with a 100x objective (CFI Apo TIRF 100X Oil, Nikon) and a 647 nm laser (laser power set to 100% and exposure to 40 ms). The acquired images were processed and converted into .txt files with Nikon NIS elements software. A MATLAB GUI previously developed in the lab (Truong Quang et al., 2021) was used to extract cortex thickness. In short, dSTORM images of isolated blebs were converted into intensity heatmaps and line-scans were traced across the cortex along the bleb contour. Full width at half maximum (FWHM) for each line-scan was then measured and averaged. Custom MATLAB scripts were used to segment blebs and calculate the bleb radii (extracted from the bleb curvature for circular blebs; for elongated blebs the cross-sectional area was computed and an equivalent radius was extracted). Merged membrane and cortex images were obtained by overlaying membrane confocal images and STORM reconstructed cortex images.

### Cryo-ET and tomogram reconstruction

For cryo-ET, 5.5 µL of isolated bleb suspension was mixed with 1.5 μL of fiducial gold markers (10 µm diameter beads, diluted 1:500, Aurion). This bleb/beads suspension was then applied onto glow discharged plasma-cleaned carbon-coated copper grids (GR1/2, Quantifoil) and manually blotted for ∼3 seconds with 55 mm Whatman filter paper (ThermoFisher) to obtain a thin film. The majority of blebs were lost during the blotting, but sufficient blebs remained for cryoET imaging. Grids were plunge-frozen in liquid ethane. A FEI Titan Krios transmission electron microscope was used to acquire isolated blebs tilt-series. The tilt-series angular range covered from −60 to +60 with 2° increments. The magnification used was 42.000x in super resolution mode, resulting in a pixel size of 1.34 nm. Defocus and electron dose were set to −4 and 7.5 e^-^/pixel/sec, respectively. Image stacks were drift corrected using a built-in Digital Micrograph module. MATLAB TOM toolbox was used to perform fiducial gold markers alignment and tomograms back-projections reconstruction (Nickell et al., 2005).

### Filaments segmentation, data analysis and density heatmaps

Amira and Amira XTracing module (FEI Visualization Sciences Group) were used to process tomograms and perform actin filaments segmentation. Tomograms were filtered with a Non-Local Means Filter in the XY plane. The filtering step was performed with a Quadro K6000 CUDA device. DPSV denoising filter (Rusu et al., 2012), Nonlinear Anisotropic Diffusion filter (Frangakis and Hegerl, 2001) and other Amira filters were tested but Non-Local Means Filter allowed optimal noise reduction. Filter search window was set at 21 and similarity value to 0.6.

Filament segmentation was carried on following (Rigort et al., 2012). Briefly, actin filaments were modeled as cylinders with a diameter of approximately 7 nm. First, denoised tomograms are traversed by a template cylinder of appropriate diameter and 54 nm in length, which is locally fitted to the data and used as a seed. A measure of similarity is used to compute the local correlation between voxels. The cross-correlation maps are then processed with a tracing algorithm, where the likelihood that two neighboring voxels belong to the same filament is calculated. Parameters chosen were similar to (Rigort et al., 2012); since filaments do not have a hollow axial section, the “inner cylinder radius” parameter was set to zero. Segmented tomograms were subjected to a manual inspection to remove false positives. For example, thicker regions of the plasma membrane and shadows of the fiducial gold beads could occasionally be detected as filaments. Inspected filaments coordinates were then exported as a .txt file. Data mining and relational databases were used to extract relevant geometrical information from filament datasets. Computations were performed with custom MATLAB and Python scripts, except extraction of the segmented bleb volume (used to estimate the volume fraction covered by actin filaments), which was done using the Amira software. Filament thickness in Figure 2C was measured as the FWHM of the intensity profile in the processed tomograms. Straightness was defined as linear length (linear distance between filament starting and ending point) divided by contour length. Custom Python scripts were used to identify branching and crossing points. A selected threshold distance defined the size of the search region for branching or crossing points. A crossing point was recorded when the distance between bodies of two filaments was smaller than the search region, defined as the cross-linking distance considered ± 3 nm. Density heatmaps of actin filaments in blebs were obtained by partitioning the tomogram volume into XY bins (spanning the entire z height of the tomogram), and computing the filaments density (total length of filamentous actin in a given bin). Bin size was kept constant for all the blebs considered. Heatmaps computation was performed in Python.

### Statistical Analysis

Student t-test, Welch’s t-test and D’Agostino-Pearson normality tests were performed with Prism software (GraphPad). Given the difference in sample size in the comparison of cortical thicknesses between isolated blebs and entire cells, we performed bootstrapping for hypothesis testing, using a custom MATLAB script; the number of iterations adopted was 10000.

